# SynaptopathyDB: a resource for studying the genetic and synaptic basis of nervous system disorders

**DOI:** 10.1101/2025.06.27.661932

**Authors:** Oksana Sorokina, Digin Dominic, Àlex Bayés, J. Douglas Armstrong, Seth G.N. Grant

## Abstract

Synaptic dysfunction resulting from pathogenic variants in genes encoding synaptic proteins is a major contributor to brain and behavioural disorders, collectively termed synaptopathies. To facilitate research into the genetic basis and clinical manifestations of synaptopathy we have created SynaptopathyDB, an online resource that integrates data from 64 mammalian synapse proteomic studies and multiple genetic and phenotypic resources. We identified a consensus set of 3,437 mammalian synapse proteins from presynaptic and postsynaptic compartments, which have wide application in genetic and omic studies. Mutations in 954 genes encoding 28% of the consensus synapse proteome were associated with 1,266 OMIM diseases of the central and peripheral nervous system. We present findings that underscore the pervasive role of synaptic gene variants in the phenotypes of neurological, psychiatric, developmental, and systemic disorders highlighting the significant burden they impose on individuals and healthcare systems. SynaptopathyDB is a versatile platform and discovery tool for understanding the role of synapse proteins and genetic variants in human disease phenotypes.

## Introduction

Synapse proteome datasets have had an immense impact on our understanding of human brain disease. The realisation that mutations in genes coding for synaptic proteins cause human brain disorders arose from proteomic studies of the postsynaptic proteome of excitatory synapses in 2000, which linked synaptic proteins to three genetic disorders causing severe neurological symptoms^1^. By 2011, mutations in 199 genes encoding postsynaptic proteins had been linked to 133 nervous system disorders^2^. In addition to revealing causal mechanisms for many rare monogenic disorders, the synapse proteome played a critical role in unravelling the biological mechanisms of polygenic disorders, including schizophrenia^3-8^, which is now recognised as a synaptic disorder. Recent large-scale genetic studies of depression have leveraged synapse proteome data to reveal that, like schizophrenia, the disorder arises from genetic variants targeting many diverse postsynaptic proteins^7,9-12^. In addition to revealing the mechanisms of common psychiatric disorders, the synapse proteome datasets have been instrumental in understanding the biological basis of the genetic variation underlying IQ, which also shows an enrichment in genes encoding synaptic proteins^13-15^.

Since 2011, there have been significant advances in the depth, scale and throughput of human genome sequencing, leading to the discovery of many more disease associated mutations. Concurrently, numerous proteomic studies have mapped the protein composition of mammalian synapses, expanding the catalogue of known synaptic proteins^16^. However, there are no centralised and readily searchable resources that integrate data on synaptic proteomes, genetics, diseases, and phenotypes. The development of such a resource would greatly facilitate both simple and complex analyses. At present, it is not straightforward to retrieve detailed associations between proteins involved in synaptopathies and their associated diseases or phenotypes. Generating consensus sets of synaptic proteins across multiple studies would greatly assist genetic association and gene enrichment analyses by providing a common reference. A comprehensive and accessible resource that would support systematic investigations into the relationships between synaptic proteins and phenotypes is lacking.

Here, we present SynaptopathyDB (www.synaptopathyDB.org), a comprehensive platform designed to facilitate exploration of the synaptic basis of brain and peripheral nervous system disorders via three intuitive search interfaces: Gene Search, Disease Search, and Publication Search. Users can efficiently retrieve relevant information using autocomplete suggestions, flexible query terms and refine results with filters such as subsynaptic protein localization. Expandable result panels provide detailed annotations and direct links to external databases including Online Mendelian Inheritance in Man (OMIM)^17,18^, International Classification of Diseases (ICD-11)^19^, Human Phenotype Ontology (HPO)^20^, Gene Ontology (GO)^21^, Synaptic Gene Ontology (SynGO)^22^, UniProt^23^ and PubMed. In addition to search functionalities, SynaptopathyDB offers interactive visualizations for tracking disease associations, discovery rates, and publication trends over time through customizable plots. For advanced users the platform provides an application programming interface (API), enabling programmatic access to raw data and complex queries. Together, these features position SynaptopathyDB as both a reference database and an integrated research tool for connecting synaptic proteins, genetic variation, and human disease, providing a new tool that will help development of treatments for synaptopathies.

### Identifying a consensus set of synapse proteins

We previously reported a database comprising 58 proteomic studies characterising the protein composition of mammalian synaptic fractions^16^. This has now been expanded to include 64 studies, collectively reporting over 8,000 unique proteins^24^. These investigations have primarily focused on human and rodent models, whose synaptic proteomes are known to be highly conserved across species^2,25-28^. However, many of the proteins identified in these studies have been reported only once and might represent experimental artefacts or contaminants. To improve data confidence, we defined a highly-reproducible consensus set of 3,437 synaptic proteins, each supported by evidence from at least five independent studies (see methods). This curated set is hereafter referred to as SynProteome^CONS^.

The synapse comprises two principal substructures: the presynaptic and postsynaptic terminals, which are interconnected by proteins within the synaptic cleft. The presynaptic bouton contains synaptic vesicles loaded with neurotransmitters, while the postsynaptic terminal houses multiprotein complexes that anchor neurotransmitter receptors, cell adhesion molecules, scaffolding proteins, and signalling molecules to the cytoskeleton. The proteomes of synaptosomes — isolated whole synapses — as well as presynaptic and postsynaptic compartments and their constituent complexes and substructures have been characterized in the 64 proteomic studies. To produce consensus sets of proteins for the presynaptic and postsynaptic terminals within SynProteome^CONS^ we compared the proteins characterised by the different methodologies and found eight groups (Supplementary Figure 1A), which we reduced to three (Supplementary Figure 1B): SynProteome^POST^ containing 1,354 proteins only found in postsynaptic preparations; SynProteome^PRE+^ comprising 456 presynaptic proteins plus proteins found in both presynaptic and postsynaptic fractions; and SynProteome^SYN^ comprising 1,627 proteins isolated from synaptosomes but not specifically allocated to presynaptic or postsynaptic compartments.

**Figure 1.**
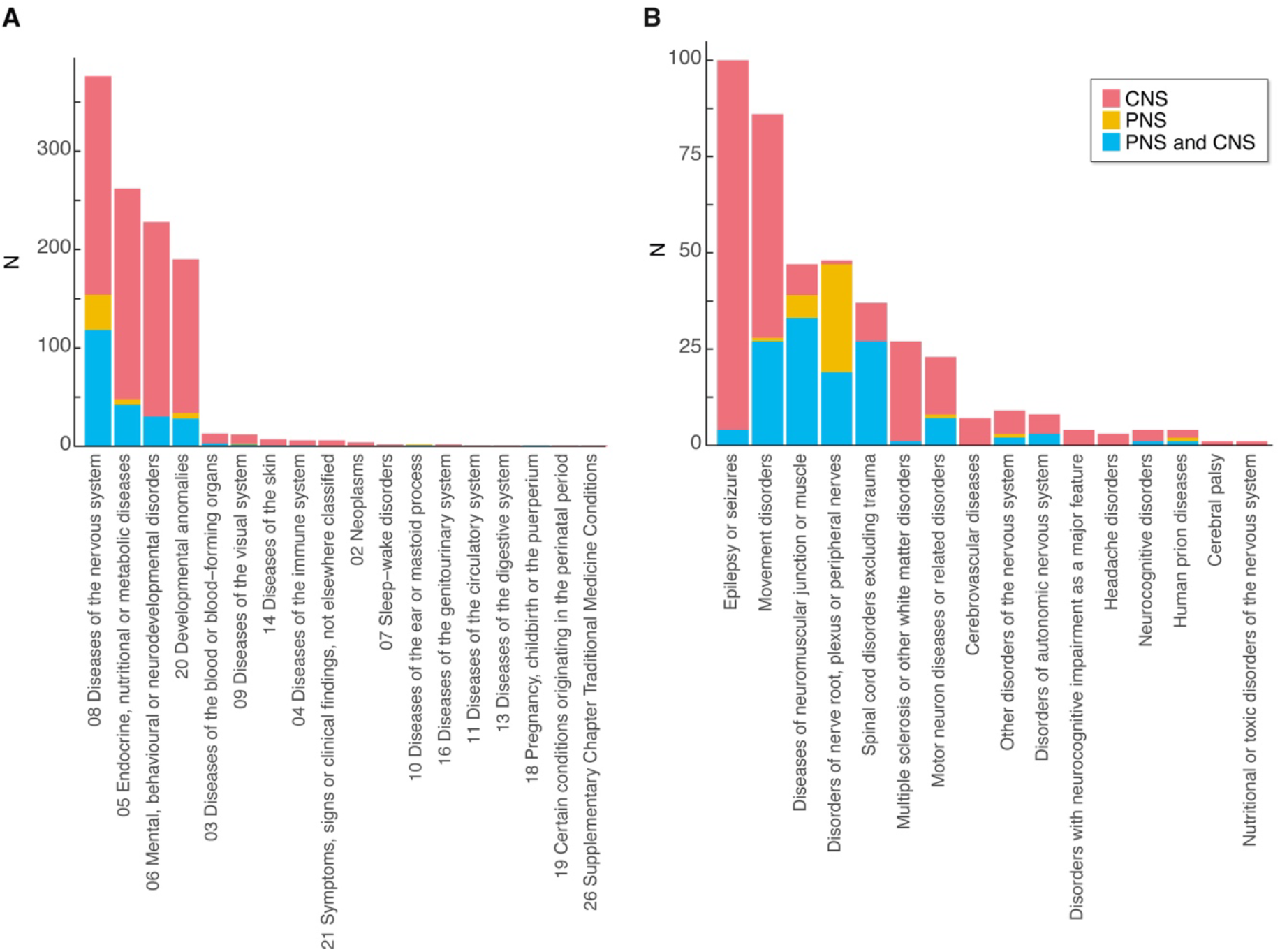
Diseases associated with synaptic gene mutations. A. The number of OMIM-associated genes associated with each ICD-11 chapter subdivided into those associated with CNS, PNS, or both, and their phenotypes. B. The number of OMIM-associated genes associated with disorders within ICD-11 Chapter 8 (Diseases of the nervous system).

### Disease-relevant genetic variants target all synapse compartments

To identify disease-relevant variants targeting synaptic proteins, we cross-referenced each of the 3,437 SynProteome^CONS^ genes with the OMIM database^17,18^, revealing 1,236 (35.9%) genes with variants associated with one or more diseases (Supplementary Table S1). A high percentage of genes in each compartment were targeted by mutations with OMIM phenotypes: 45% (206/456) of SynProteome^PRE+^, 38% (516/1359) of SynProteome^POST^ and 32% (514/1627) of SynProteome^SYN^ genes (Table 1).

**Table 1.** Synapse proteome subsets and their associated disease genes. The number of genes (Unique genes) in each proteome set is shown. For each proteome set the number of genes encoding that set (Unique genes) and the number with OMIM IDs (Unique genes with OMIM ID). Additional columns show the number and percentage of disease genes within each proteome set associated with phenotypes with CNS, PNS, combined CNS and PNS. Total NS column, combining the PNS, CNS and PNS&CNS, corresponds to the number of OMIM diseases that have any of the nervous system phenotypes. Genes associated with non-nervous system phenotypes (nonNS) are shown.

| Proteome set | Unique genes | Unique genes with OMIM ID | CNS | PNS | PNS&CNS | Total NS | nonNS |
| --- | --- | --- | --- | --- | --- | --- | --- |
| SynProteome<sup>CONS</sup> | 3437 | 1236 | 709 (57%) | 25 (2%) | 220 (17.8%) | 954 (78%) | 282 (22.8%) |
| SynProteome<sup>POST</sup> | 1354 | 516 | 296 (57.3%) | 15 (2.9%) | 93 (18%) | 404 (78.2%) | 112 (21.7%) |
| SynProteome<sup>PRE+</sup> | 456 | 206 | 129 (62%) | 2 (1%) | 43 (20.1) | 174(84.4%) | 32 (15%) |
| SynProteome<sup>SYN</sup> | 1627 | 514 | 284 (55.2) | 8 (1.5%) | 84 (16.3) | 376 (73.1) | 138 (26.8%) |

We next asked how many of these disease-relevant genes were linked to diseases that involve the central (CNS) and peripheral (PNS) nervous system or to non-nervous system disorders (Table 1). Seventy-eight percent (954 genes) of SynProteome^CONS^ disease-relevant genes have either a CNS or PNS phenotype, with similar percentages in each of the synaptic compartments. Of the different synaptic compartments, SynProteome^PRE+^ showed the highest levels of enrichment (total nervous system, 84.4%, 174 genes, p = 1.4 × 10^−7^; CNS-associated, 62%, 129 genes, p = 1.7 × 10^−5^, hypergeometric test).

### Types of brain diseases and their phenotypes

To further understand the types of brain disorders caused by synaptic mutations we integrated SynProteome^CONS^ with ICD-11^19^. ICD-11 organizes disorders into numbered chapters, several of which are directly relevant to the nervous system, including Chapter 6 (Mental, behavioural or neurodevelopmental disorders) and Chapter 8 (Diseases of the nervous system). As shown in Figure 1A, ranking the 20 ICD-11 chapters by the number of associated synaptic gene variants revealed that the majority of these mutations are concentrated in four chapters. The chapter with the highest number of associated genes is Chapter 8, comprising 343 of the 1,236 genes, with Chapter 6 also prominently represented (227 genes). A substantial number (227 genes) of disease-associated synaptic genes were also mapped to Chapter 20 (Developmental anomalies), which includes disorders affecting brain structure and nervous system formation, and (316 genes) to Chapter 5 (Endocrine, nutritional or metabolic diseases), which includes metabolic dysfunctions — such as impaired energy metabolism — that often compromise neuronal function and may manifest with diverse nervous system phenotypes. Plotting the contribution of CNS and PNS diseases to each of these chapters shows the major contribution of synaptic pathogenic variants to the CNS (Fig. 1A).

We also asked if any of these disease categories are more closely associated with any specific synapse compartments (Table 2). We found that SynProteome^POST^ shows a significant enrichment in genes associated with mental/neurodevelopmental disorders (Chapter 6) (p = 7.9 × 10^−6^) and that SynProteome^PRE+^ shows significant enrichment for nervous system diseases (Chapter 8, p = 2.4 x 10^−8^) and modest significance in developmental disorders (Chapter 20) (p = 0.049). Next, we examined the subcategories within diseases of the nervous system (Chapter 8), which describes many classes of disorders that traditionally fall under the large family of neurological diseases (Fig. 1B). Epilepsies and movement disorders contain the highest number of synapse gene variants and both are dominated by CNS-acting gene mutations.

**Table 2.** Synapse proteome subsets and their associated disorders. The number of genes (Unique genes) in each proteome set is shown. For each proteome set the number of genes encoding that set (Unique genes) and the number with OMIM IDs (Unique genes with OMIM ID). Additional columns show the number and percentage of disease genes within in ICD-11 chapters 5, 6, 8 and 20.

| Proteome set | Unique genes | Unique genes with OMIM ID | Chapter 8<br>Disease Of Nervous System | Chapter 6<br>Mental, Behavioural Or neurodegenerative | Chapter 5<br>Endocrine, Nutritional Metabolic | Chapter 20<br>Developmental Anomalies |
| --- | --- | --- | --- | --- | --- | --- |
| SynProteome<sup>CONS</sup> | 3437 | 1236 | 343 (28%) | 227 (18.3%) | 316 (25.5%) | 227 (18.4%) |
| SynProteome<sup>POST</sup> | 1354 | 516 | 153 (27.5%) | 121 (23.4%) | 96 (18.6%) | 100 (19.3%) |
| SynProteome<sup>PRE+</sup> | 456 | 206 | 81 (33.9%) | 44 (21.3%) | 41 (19.9%) | 39 (18.9%) |
| SynProteome<sup>SYN</sup> | 1627 | 514 | 109 (21.1%) | 62 (11.9%) | 179 (33.4%) | 88 (17.1%) |

Diseases are classified from constellations of phenotypes, which clinically are referred to as symptoms and signs. To identify the human phenotypes caused by pathogenic variants in genes encoding synaptic proteins, we mapped SynProteome^CONS^ genes involved with disease to the Human Phenotype Ontology (HPO)^20^, identifying a total of 3,986 phenotypes (Supplementary Table S2). Next, using gene set enrichment analysis we identified the phenotypes most relevant for the SynProteome^CONS^ and found 739 significantly enriched terms (hypergeometric test). The major enriched phenotype was Abnormality of the Nervous System and within this two other terms Abnormal Nervous System Physiology and Abnormal Nervous System Morphology, were also highly enriched. Figure 2A show 18 terms from these two categories which are highly significantly enriched (P< 10^−5^ – 10^−45^) in SynProteome^CONS^, SynProteome^PRE+^ and SynProteome^POST^genes. Cognitive, motor and epileptic phenotypes are highly enriched within the Abnormal Nervous System Physiology category (Fig. 2B).

**Figure 2.**
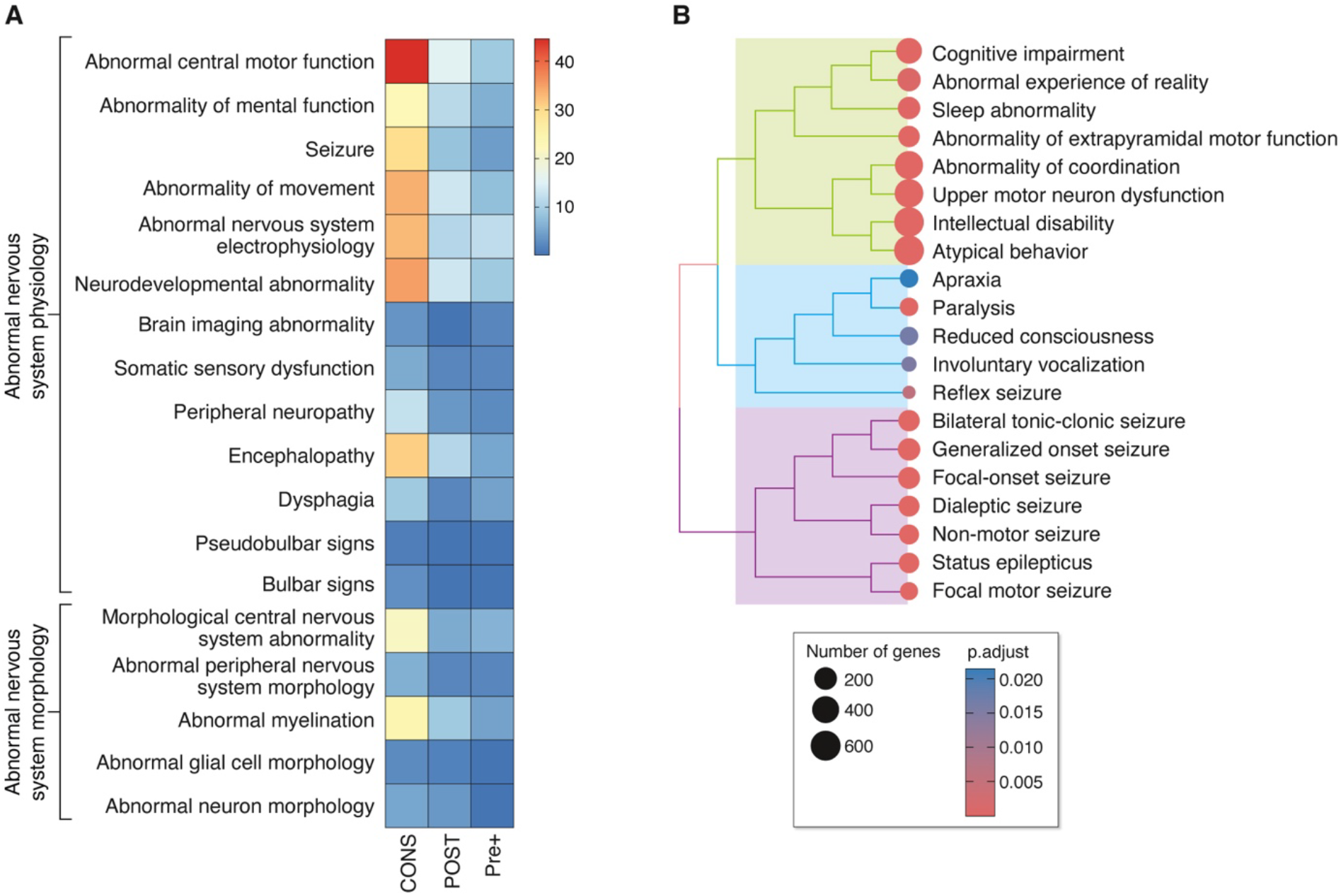
Human phenotypes of synaptic mutations. A. Highly enriched HPO phenotypes within Abnormalities of the Nervous System phenotypes. Key, log(p.adj) enrichment value. CONS, SynProteome^CONS^; POST, SynProteome^POST^; Pre+, SynProteome^PRE+^. B. Clustering of highly enriched cognitive, motor and epileptic phenotypes within the Abnormal Nervous System Physiology category.

### Declining rate of discovery of diseases linked to synapse mutations

How many more diseases arising from genetic variants in genes coding for synaptic proteins remain to be discovered? To evaluate this, we built a display in SynaptopathyDB that produces a chronology of the number of diseases linked to the 1,236 genes (Figure 3A). This shows that the total (nervous and non-nervous system) number of diseases discovered per year reached a peak in 2013 and since 2017 has been steadily declining. This trend is reflected in the cumulative disease discovery plot shown in Figure 3B. These data show that 1,768 (1,266 nervous and 502 non-nervous system) diseases were identified by 2024, suggesting that the spectrum of diseases and phenotypes associated with synaptic gene variants for this set of genes is reaching completion. At present, 64% of SynProteome^CONS^ is not associated with disease variants and it seems plausible that many genes encoding this subset will be candidates for future discovery.

**Figure 3.**
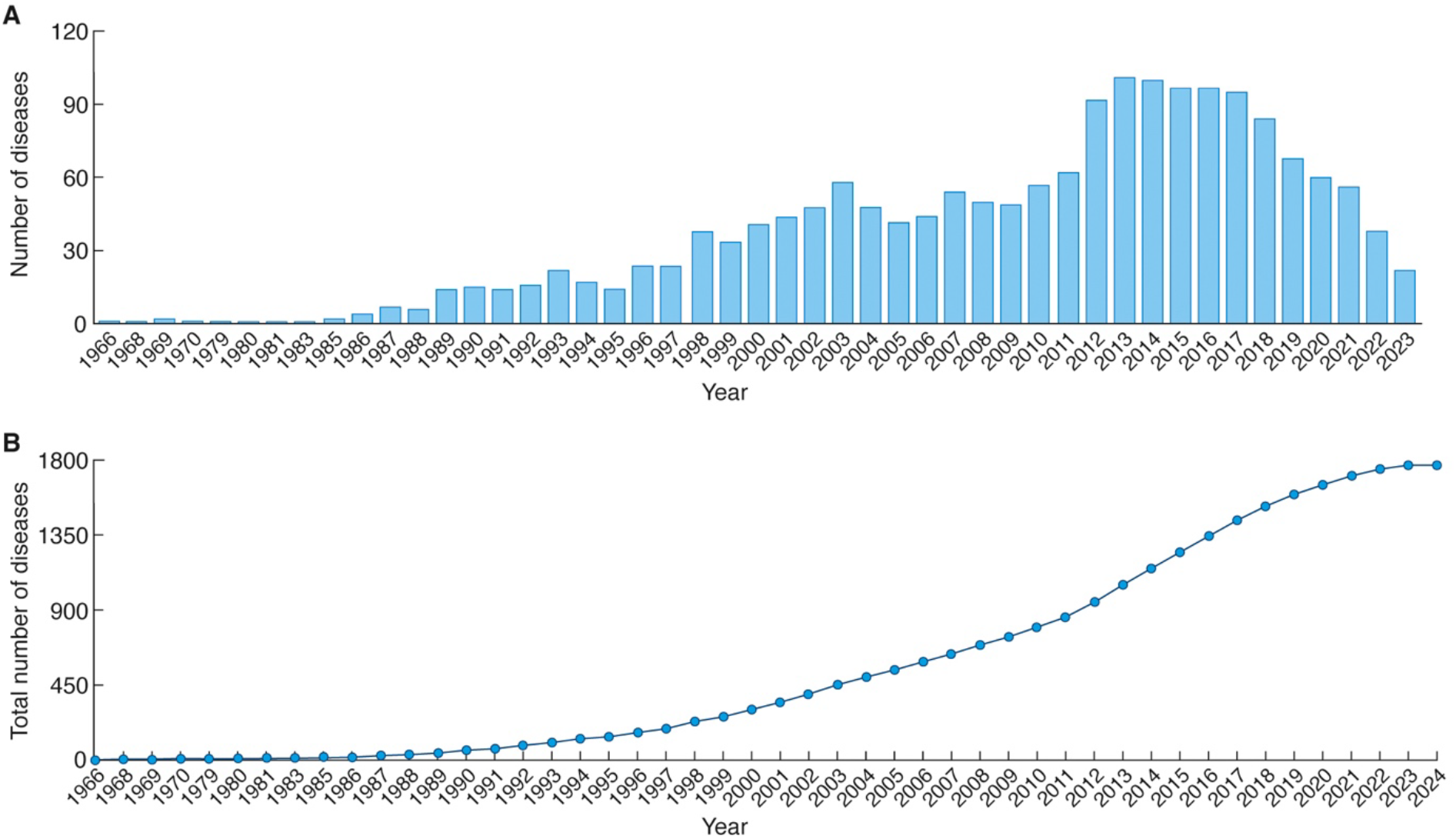
Declining rate of discovery of diseases linked to synapse mutations. A. The number of diseases discovered each year that were later reported to be linked to synaptic gene mutations B. The cumulative number of diseases discovered per year.

### Using the SynaptopathyDB website

The website features a comprehensive search system with three main entry points: ‘Gene Search’, which allows users to identify which nervous system disorders are caused by a given gene or set of genes; ‘Disease Search’, which allows users to perform the opposite search, identifying which synaptic genes are associated with specific disorders; and ‘Publication Search’, which retrieves the scientific publications with gene-disease connections. The platform supports quick search with autocomplete^29-31^, multiple identifier queries, and highlighted results. Filtering options enable users to refine searches based on protein localization within the synapse or specific gene mentions in publications. Expandable panels, link-outs to external databases, and rich metadata, including subcellular locations and mutation details enhance utility. Data visualization tools provide interactive disease trend analyses, track discovery rates over time, and display annual mentions of disease-associated genes through various chart types. Additionally, the platform offers API endpoints for exporting structured datasets, including synapse proteome lists, in CSV format for further analysis.

With the expected discovery of further mutations targeting synaptic proteins we plan regular updates of SynaptopathyDB and ongoing expansion of functionality that will assist in classifying and understanding the synaptic basis of brain disease. The consensus proteome lists described herein have wide application. For example, they can be applied to large-scale genetic (e.g. GWAS, CNV) studies, transcriptomic and proteomic studies of tissue and cell types, genetic screens and drug discovery. Our analyses also underscore the importance of the molecular complexity of the synapse proteome, and its genetic underpinnings, in controlling the diverse and complex behavioural repertoire of humans.

## Methods

### Synaptic proteome database

Synaptic studies for presynaptic, postsynaptic, synaptic vesicle and whole synaptosome compartment were identified by manual curation of PubMed, starting from year 2000, which resulted in 64 studies^24^. Protein/gene lists from the published synaptic proteome studies were combined into the total list of synaptic components. Identifiers from each study were mapped to stable IDs including: Entrez Human and Mouse, Uniprot and MGI IDs. Each protein is associated with literature ID and curated list of metadata such as species (mouse, human, rat), respective synaptic compartment, brain, experimental methods, Gene Ontology (GO) and disease associations (see below). The synaptic proteome datasets are available in SQLite format^24^.

### Building the consensus synaptic protein list

Using the synapse proteome database^24^ we have selected the human genes that were identified at least five times in any of the following synaptic compartments: postsynaptic, presynaptic, whole synaptosome, synaptic vesicle. We assume that, given threshold guarantees, the specific protein is highly likely to be associated with this compartment, and excluded the proteins with a few identifications across multiple compartments as potential contaminants.

### Disease annotation

We used gene-disease annotation data collected from the OMIM^17^, GeneRIF^32^and Ensembl variation^33^ databases. Using the topOnto package, the annotation data were standardised using MetaMap^34^ and NCBO Annotator ^35,36^ to recognize terms found in the Human Disease Ontology (HDO)^37,38^. Recognised disease ontology terms were then associated with gene identifiers and stored locally.

For each of the consensus genes we extracted gene and phenotype entries from OMIM using the mim2gene file linking via their Entrez Gene identifiers. Diseases associated with SynProteome^CONS^ proteins were further classified using information in the OMIM entry (Supplementary Table S1), into CNS, PNS, both, or other. All diseases were then grouped into chapters of the International Classification of Disease (ICD-11) developed by the World Health Organisation (WHO). Those OMIM diseases that could not readily be found in ICD-11 were assigned to one of the ICD-11 chapters based on their symptoms, if possible. Diseases in Chapter 8 (Diseases of the nervous system) were divided into more specific blocks within that chapter.

### Human Phenotype Ontology gene set enrichment analysis

Gene lists representing human illnesses and conditions were downloaded from the Human Phenotype Ontology (HPO). The HPO database curates OMIM diseases and genes into a hierarchical format with more higher-level terms representing more general clusters of phenotypes. Genes linked to each condition are also documented in the HPO. This dataset was downloaded on January 20, 2025. Enrichment of phenotype genes in the SynProteome^CONS^ relative to the genome was assessed on phenotypes that had more than 20 genes in the genome. Significances of overrepresentation of disease conditions in the SynProteome^CONS^ were assessed by hypergeometric statistics corrected by the Benjamini-Hochberg false discovery rate, implemented as *fora* function in *gsea* R package. For comparison, a non-synaptic protein list was built by extraction of our latest total synaptic list (10,500 genes, not shown) from whole human genome (NCBI database) downloaded in January 2025, which resulted in 9,667 genes. Overrepresentation of each phenotype relative to the genome was computed separately for synaptic (SynProteome^CONS^) and for the non-synaptic gene lists.

### Web-based resource

SynaptopathyDB is implemented as a comprehensive web application with a user-friendly interface designed to facilitate exploration of synaptic protein-disease relationships. The platform provides the following primary search interfaces:

### Unified search

The homepage features a unified search functionality that allows users to simultaneously search across genes, diseases, and publications. The autocomplete feature provides real-time suggestions, supporting multiple identifier types including:

- Gene names and identifiers (Human/Mouse/Rat Entrez, MGI, Ensembl ID, Alias)^29-31^.
- Disease names and ontology identifiers
- Publication details (PMID, title, year)

### Gene search

The gene search interface enables comprehensive exploration of individual genes, providing:

- Detailed gene information including multiple species identifiers^29-31^
- Associated publications
- Disease associations
- Mutation details
- Localization within synaptic compartments
- Filtering options by consensus status
- Downloadable data in CSV format

### Disease search

The disease search interface allows users to:

- Explore diseases and their associated synaptic genes
- View detailed mutation information
- Filter genes by consensus status
- Access publication links
- Generate downloadable reports

### Publication search

The publication search feature provides:

- Comprehensive details of synapse proteomics studies
- Filtering of associated genes and mutations
- Localization and method information
- Interactive pagination and export capabilities

### Data visualization and analysis tools

SynaptopathyDB offers several chart visualizations:

- Disease classification trends
- Protein localization history
- Discovery rate analyses
- Compartment-specific protein distributions

### Technical infrastructure

- Frontend: React.js with Tailwind CSS
- Backend: Flask (Python)
- Database: SQLite
- Interactive components powered by Recharts and other visualization libraries

### API and data access

The platform provides RESTful API endpoints for:

- Programmatic data retrieval
- CSV export of comprehensive datasets
- Flexible querying across genes, diseases, and publications

The web application is designed to be intuitive, responsive, and accessible across different devices, supporting researchers in exploring the complex relationships between synaptic proteins, genetic variations, and human diseases.

## Acknowledgments

OS, JDA and SGNG were supported by BBSRC Grant number BB/X009343/1, ‘Development of a computational model of synaptome architecture’. DD is supported by a grant from The Wellcome Trust [218293/Z/19/Z] to SGNG. AB for PID2021-124411OB-I00 (MINECO/MCI/AEI/FEDER, EU), AGAUR (2021 SGR 01005) and the CERCA Programme/Generalitat de Catalunya. P. Alchonel Albeida, literature annotation. D. Maizels, artwork. C. Davey, editing.

## Data availability

For the purpose of open access, the author has applied a CC-BY public copyright licence to any Author Accepted Manuscript version arising from this submission. Synapse proteome data are available from Edinburgh DataShare at https://doi.org/10.7488/ds/3771

## Supplementary Table legends

**Supplementary Table S1**. List of 3,437 consensus synaptic genes with respective OMIM and ICD-11 phenotypes.

**Supplementary Table S2**. HPO phenotypes identified for consensus synaptic genes.

---

**Supplementary Figure S1.**
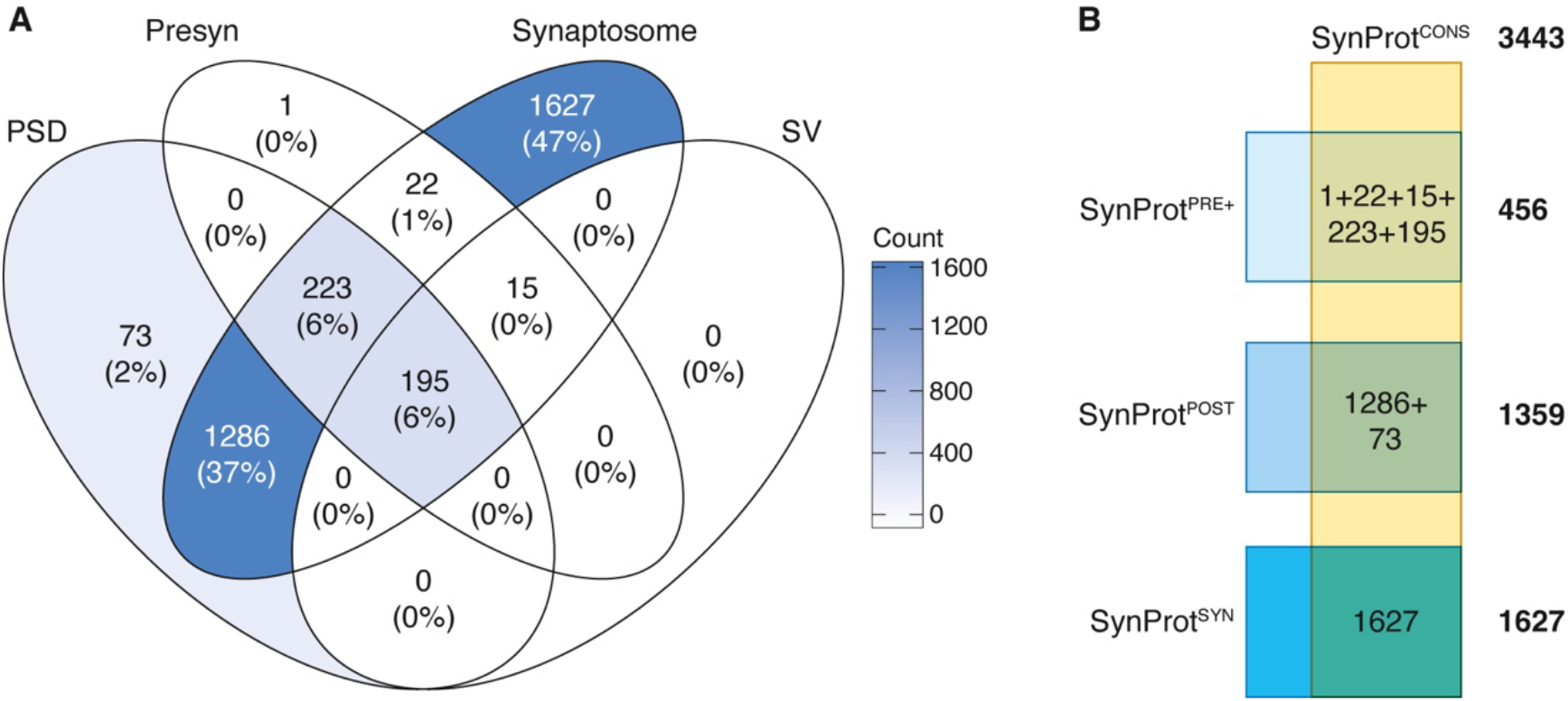
Defining consensus synapse proteome subsets. A. Venn diagram showing numbers of proteins in different synaptic fractions from multiple studies. Presyn, presynaptic; PSD, postsynaptic density; SV, synaptic versicle. B.Defining the SynProteome^CONS^, SynProteome^POST^, SynProteome^SYN^ and SynProteome^PRE+^ sets from groups in A.

